# After-effects of Parieto-occipital Gamma Transcranial Alternating Current Stimulation on Behavioral Performance and Neural Activity in Visuo-spatial Attention Task

**DOI:** 10.1101/2024.10.06.616926

**Authors:** Tianyi Zheng, Yunshan Huang, Masato Sugino, Kenta Shimba, Yasuhiko Jimbo, Kiyoshi Kotani

## Abstract

Visuo-spatial attention enables selective focus on spatial locations while ignoring irrelevant stimuli, involving both endogenous and exogenous attention. Recent advancements in transcranial alternating current stimulation (tACS) have shown promise in modulating these attentional processes by targeting electrical oscillations in specific brain areas. Despite evidence of online effects of tACS on visuo-spatial attention performance, whether tACS can produce lasting after-effects on behavioral performance and neural activity remains unknown. This study explored these after-effects using a single-blind, sham-controlled, between-group design. Eighteen young healthy participants were equally divided into two groups receiving either sham or active gamma tACS at 40 Hz targeted at the right parieto-occipital region. Each participant performed a version of the Posner cueing task with EEG recording before and after the tACS intervention. The active tACS group exhibited greater reductions in reaction time compared to the sham group. These changes were not uniform across different attention types, suggesting specific enhancements in cognitive processing. EEG analyses revealed trial-type-specific modulation of event-related potentials, including amplitude and latency of N1 and P3 components, that paralleled the behavioral effects. Additionally, frequency-specific changes in oscillatory power during the cue-target interval—decreased alpha power and increased gamma power—as well as reduced long-range temporal correlations were observed more broadly across conditions. While limited by a small sample size, these preliminary findings provide convergent behavioral and electrophysiological evidence that parieto-occipital gamma tACS can induce lasting, condition-specific after-effects on visuo-spatial attentional networks.

## INTRODUCTION

Visuo-spatial attention enables selective prioritization of visual information at specific locations while filtering irrelevant input, thereby allocating limited processing resources to behaviorally relevant stimuli (Posner et al. (1980)). A widely used framework distinguishes *endogenous* (voluntary, goal-directed) attention from *exogenous* (stimulus-driven) attention (Corbetta and Shulman (2002)). These attentional modes can be assessed with variants of the Posner cueing task (Posner et al. (1980)), in which reaction times (RTs) to a visual target differ depending on whether the preceding cue validly or invalidly predicts target location.

Converging evidence suggests that attentional orienting and target processing rely on coordinated dynamics within parietal and occipital regions, and that neuronal oscillations contribute to these dynamics across multiple frequency bands, including alpha and gamma (Klimesch (2012); Han et al. (2022)). In particular, gamma-band activity in parietal and occipital cortices has been linked to attentional selection and the integration of task-relevant sensory information (Gruber et al. (1999); Bosman et al. (2012); Magazzini and Singh (2018)). Because endogenous and exogenous attention differ in their reliance on top-down versus bottom-up control, neuromodulation of parieto-occipital dynamics may not affect these attentional modes uniformly across cueing conditions. For example, endogenous orienting is typically more dependent on goal-driven control signals engaging dorsal parietal mechanisms, whereas exogenous orienting is more strongly driven by salient sensory events and rapid stimulus-driven capture, which may place different demands on occipital processing and attentional reorienting (Corbetta and Shulman (2002)).

Transcranial alternating current stimulation (tACS) is a non-invasive technique that delivers oscillatory electrical currents through scalp electrodes to modulate brain activity (Elyamany et al. (2021)). Gamma-frequency stimulation, often operationalized at 40 Hz, targets putative gamma-band mechanisms that are crucial for cognitive processing and sensory integration, enhancing neuronal synchronization and connectivity essential for processing complex visual stimuli and improving visuo-spatial attention (McDermott et al. (2018); Jia et al. (2011)). Moreover, 40 Hz stimulation has been shown to improve brain function through both audiovisual (Chen et al. (2022)) and tACS interventions (Hopfinger et al. (2017)), positively impacting synaptic plasticity and brain network connections. Accordingly, several studies have reported online behavioral effects of 40 Hz tACS in visuo-spatial attention tasks (Kasten et al. (2020); Hopfinger et al. (2017); Reteig et al. (2017)). Notably, Hopfinger et al. reported that 40 Hz tACS over right parietal regions during task performance modulated cueing-related behavior in a manner that depended on attention type and cue validity, supporting the relevance of this frequency for probing gamma-related mechanisms in visuospatial orienting (Hopfinger et al. (2017)). At the same time, the extent to which conventional-intensity scalp-applied tACS produces spatially specific, on-target modulation—as opposed to broader field spread, peripheral co-stimulation, and/or measurement confounds—remains an active topic of debate (Vö rö slakos et al. (2018)). This motivates careful interpretation of montage specificity and encourages convergent behavioral and electrophysiological evaluation.

Critically, most tACS studies in visuo-spatial attention have emphasized *online* effects during stimulation, whereas the presence and nature of *offline* after-effects remain less clear. Offline after-effects are theoretically important because they may reflect lasting changes in neural dynamics beyond transient entrainment. Consistent with this view, experimental work suggests that stimulation can shape event-related rhythmic activity in ways that can produce after-effects on evoked oscillatory responses, supporting broader mechanistic accounts for after-effects (Wischnewski and Schutter (2017)). However, whether parieto-occipital 40 Hz tACS induces measurable offline changes in visuo-spatial attention performance and task-evoked neural signatures remains unknown.

To address this gap, we used a single-blind, sham-controlled, between-group design in which participants performed a Posner cueing task before and after sham or active parieto-occipital gamma tACS (Posner et al. (1980)). We quantified behavioral performance using RTs across endogenous/exogenous and valid/invalid conditions. To probe stimulation-related changes in neural processing, we analyzed EEG using complementary measures capturing temporally specific and state-like dynamics: event-related potentials (ERPs), oscillatory power during the cue-target interval (CTI), and long-range temporal correlations (LRTC). For ERPs, we focused on N1 (90–200 ms) and P3 (250–400 ms) components evoked by target onset, given their established links to visuo-spatial attentional processing (Di Russo et al. (2003); Miniussi et al. (1999)) and the broad utility of ERPs for indexing time-locked cognitive processes (Sur and Sinha (2009)). Moreover, RT has been associated with both ERP amplitude and latency in prior work (Krieger and Dillbeck (1987); Kreegipuu and Allik (2007)).

We hypothesized that, relative to sham, active gamma tACS would induce offline modulation of visuo-spatial attention that may differ across cueing conditions (Corbetta and Shulman (2002)). Specifically, we predicted: (H1) behavioral performance would change from pre- to post-stimulation in the active group relative to sham, potentially in a trial-type-dependent manner; (H2) target-evoked N1/P3 component amplitude and/or latency would be modulated after active stimulation (Di Russo et al. (2003); Miniussi et al. (1999)); (H3) CTI oscillatory power, particularly in alpha and gamma bands, would be altered following active stimulation, given the established roles of alpha-band dynamics in selective attention and sensory suppression (Klimesch (2012); Foxe and Snyder (2011)) and the involvement of higher-frequency activity in attentional reorienting and network coordination (Spooner et al. (2020)); and (H4) LRTC during CTI would be modulated in association with behavioral changes, consistent with prior links between LRTC and cognitive performance (Palva et al. (2013); Meisel et al. (2017); Sugimura et al. (2021)) and evidence relating beta/gamma LRTC to sustained visual attention performance (Irrmischer et al. (2018)).

## MATERIALS & METHODS

### Participants

Subjects were 18 healthy volunteers (3 females), ages 24.8 ± 2.0 years (M ± SD). None of the subjects had any history of neurological or psychiatric problems. All subjects had normal or corrected-to-normal visual acuity and no history of psychiatric illness, neurological disorder, or incident. Since caffeine, nicotine, and alcohol would influence brain dynamics (Vergara et al. (2018)), cognitive function (Lorist and Snel (1998)), and the effect of tACS (Antal et al. (2017)), subjects were asked to not take any of them within 24 hours before the experiment.

### Procedure

This study was approved by the Research Ethics Committee at The University of Tokyo based on the Research Ethics Review Implementation Rules of The University of Tokyo (21-271, 22-432). All the data in this study were collected at The University of Tokyo, Japan. The main trial was conducted according to the guidelines of the Declaration of Helsinki. The participants provided their written informed consent to participate in this study.

Prior to the experiment, the rationale and potential risk of the experiment, the potential side effects of tACS (e.g. slight tingling, burning, heat, and itching sensation over the scalp), and their right to withdraw at any time during the study were explained to the subjects.

The entire experiment was divided into three main sessions: a pre-tACS task, active or sham tACS, and a post-tACS task. We recorded the EEG signal of subjects during both pre-tACS task and post-tACS task sessions (Figure 1A). Subjects answered two questionnaires: one before the entire experiment and the other between the tACS session and the post-tACS task. The questionnaire before the entire experiment is to obtain information about age, sex, medical history, and ongoing medication. We also collected the input volume of caffeine, nicotine, and alcohol in the past month. The questionnaire answered after the tACS session is to report any tACS-induced sensations including itching, pain, burning, heat, metallic taste, fatigue, and general state.

**Figure 1.**
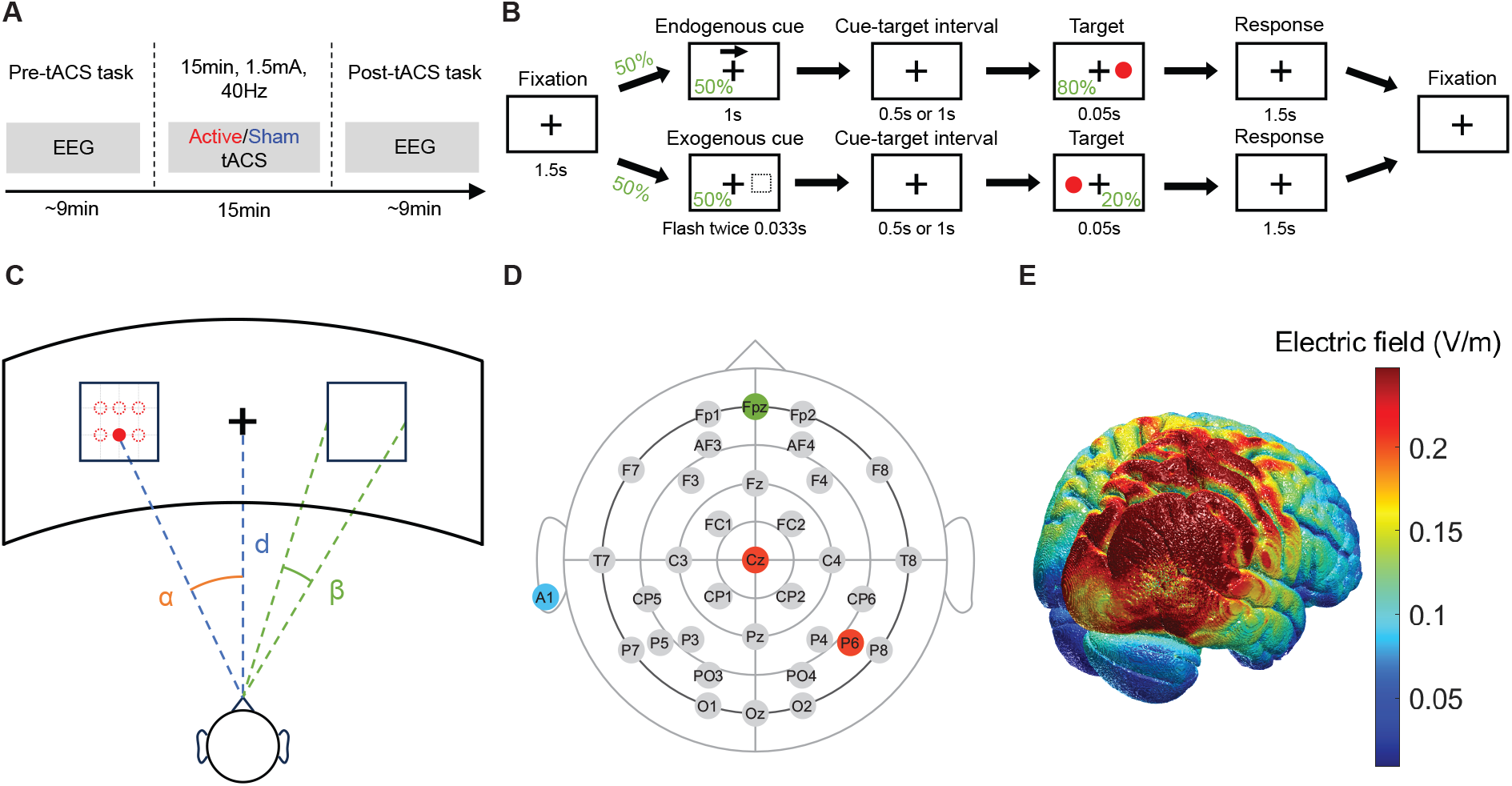
Experiment protocol. (A) Overview of experimental design. (B) Visuo-spatial attention task. (C) Layout of the display and subject in the experiment. (D) Electrode map for EEG. Ground electrode at ‘Fpz’, reference electrode at ‘A1’. ‘Cz’ and ‘P6’ are at the center of tACS electrodes. (E) Simulation of the electric field induced by the gamma tACS over the right parieto-occipital region.

Subjects sat on a comfortable chair throughout the entire experiment. We kept the laboratory quiet during two Posner cueing task sessions. During the process of answering questionnaires and tACS session, we kept the laboratory bright. During two Posner cueing task sessions, we kept the laboratory dim so that the subjects could easily see the visual target on the display.

### Visuo-spatial attention task

The visuo-spatial attention task in the current study is a version of the Posner Cueing task (Posner et al. (1980)). We utilized reaction time in a Posner cueing task (Figure 1B) as a key metric to assess the visual attention of the subjects. Each task contained 120 trials. Each trial began with a fixation, during which subjects were asked to focus on a cross symbol at the center of the screen. Fixation was followed by the presentation of an endogenous cue or an exogenous cue, randomly presented to the subjects in 50-50 probability. The endogenous cue was a centered arrow located above the fixation cross pointing left or right with a 50-50 probability. The exogenous cue was a flashing box on the left or right side of the display with a 50-50 probability. Subjects were instructed to covertly shift their attention towards the cued location while focusing on a white central fixation cross, without rotating their eyeballs or moving their heads. To reduce precise timing expectancies, the cue-target interval (CTI) was set to *T*_*CTI*_ =0.5 or 1 second with 50-50 probability (Hopfinger et al. (2017)). After the random interval, a target was presented for a very short duration (0.05 seconds). The target appeared in the same direction as the cue in 80% of trials, which were valid trials, and in the opposite direction of the cue in 20% of trials, which were invalid trials. Subjects were instructed to respond by push a space key on a U.S. keyboard as soon as possible to the appearance of the target. The response period lasted for 1.5 seconds before the fixation of the next trial. The timing of the subjects pressing the key was recorded. The reaction time was defined as the duration from the onset of the target on the display until the subject pressed the button.

To increase the field of view and reduce visual fatigue (Park et al. (2017)), we used a curved display for the task(Figure 1C). Visual information was presented against a black background on a 49-inch curved display (Philips Brilliance 499P9H), with a resolution of 5120 × 1440, and a refresh rate of 60 frames per second. The plus symbol at the center of the display was for fixation. Two squares on the left and right sides of the fixation were the places where targets would appear. The target was disc-shaped (RGB = 1,0,0; 2^◦^ of visual angle). The target could appear in one of six possible locations inside the rectangular on the screen in order to reduce the predictability. All locations of targets shared the same probability. The six target locations were at the intersections of two horizontal trisects and three vertical quadrisects of the square on each side. The distance and angles in the panel were as follows: *α*=30^◦^, *β* =10^◦^, d=0.58 m. The length of each trial depends on the type of trial because the cue duration and cue-target interval (CTI) differ across types (Figure 1B). For trials with endogenous cue and short CTI interval (N=30), they were 4.55 s long. For trials with endogenous cues and long CTI (N=30), they were 5.05 s long. For trials with exogenous cues and short CTI (N=30), they were 3.682 s long. For trials with exogenous cues and long CTI (N=30), they were 4.182 s long. In total, one session of the visuo-spatial attention task took 523.92 s (8 min and 43.92 s).

### Transcranial alternating current stimulation

Our montage of tACS selected ‘P6’ and ‘Cz’ of the international 10-20 system (Homan et al. (1987)) for electrode placement. The 10-20 system for electrode placement has been shown to reliably target desired cortex regions, including the parietal lobe (Herwig et al. (2003)). The location ‘P6’ is to stimulate the angular gyrus in the inferior parietal lobe, which is implicated in central mechanisms of movement perception and shifting of visual attention in the contralateral visual field (SAITO et al. (1992); Studer et al. (2014)). The ‘Cz’ electrode is positioned at the intersection of sagittal and coronal midlines, directing the major electric field to the parieto-occipital regions. Moreover, a previous study found that online gamma tACS delivered at ‘P6’ and ‘Cz’ of the 10-20 system results in shortening the reaction time of endogenous trials of the Posner cueing task (Hopfinger et al. (2017)). Our montage of gamma tACS is to apply stimulation towards the parieto-occipital region which is crucial for visuo-spatial attention (Bisley and Goldberg (2010); Somers and Sheremata (2013)).

To understand which brain regions are affected by the tACS, we simulated the electric field of tACS using the ROAST package (Huang et al. (2019); Huang (Y. and Datta)) and New York head MRI structural image (Huang et al. (2016)). The majority of the electrical field induced by tACS affected the right parietal lobe and the right occipital lobe, as illustrated in Figure 1E. The concurrent stimulation of parietal and occipital regions was chosen based on their critical involvement in visuo-spatial processing (Gruber et al. (1999)) and the documented benefits of targeting these areas with gamma tACS (Hopfinger et al. (2017); Kasten et al. (2020)). While the simulation results indicate activation in a wide range of brain regions, including the temporal lobe and precentral gyrus, the primary focus remains on the parieto-occipital region. The broad activation can be attributed to the nature of tACS, which often affects multiple interconnected regions due to the diffuse nature of electric fields. However, the montage’s main target remains in the parieto-occipital area, which aligns with our study’s objectives.

Before the electrical stimulation, we applied skin preparation gel (skinPure, NIHON KOHDEN, Tokyo, Japan) to targeted areas of the subjects’ scalp to reduce skin impedance. tACS was delivered by a battery-driven current stimulator (nurostym tES™, Neuro Device Group S.A., Warsaw, Poland) through two rubber carbon electrodes (Amrex CM-A102). Conductive paste (Ten20 paste, Weaver and Company, Aurora, CO) was applied to the electrodes to enable the electrodes to remain in place while allowing the transmittance of electrical signals. Two tACS electrodes (3 × 3 cm) were affixed to the location of ‘P6’ and ‘Cz’ of the 10-20 system using tape and supporting bandage. The impedance was kept below 10 kΩ during the stimulation session as monitored in real-time by the stimulator. The peak-to-peak amplitude and frequency of tACS were set to 1.5 mA and 40 Hz. Since we intended to enhance the gamma band synchronization, subjects received in-phase 40 Hz tACS, that phase difference was 0^◦^ between ‘P6’ and ‘Cz’, for the tACS intervention (ten Oever et al. (2016); Abellaneda-Pérez et al. (2020)). The active stimulation included a fade-in and fade-out of 10 seconds each and a main stage duration of 15 minutes. The sham stimulation ramped up to 1.5 mA and ramped down to 0 in 10 seconds of both the fade-in and fade-out stages. In the main stage, the sham stimulation delivered brief pulses of 2 cycles of tACS at 0.6 mA, 40 Hz every 500 ms.

Current study was sham-controlled and single-blinded which enables us to eliminate the placebo effect of stimulation (Romanella et al. (2023); Braga et al. (2021)). Subjects were randomly assigned to either the sham stimulation group (N=9) or the active stimulation group (N=9) and were unaware of which group they belonged to. To assess blinding success, a blinding survey was completed at the end of the tACS session by asking subjects to guess whether they received active or sham stimulation. The information of reported group and conditioned group were collected in Table 1. We observed that subjects were unable to guess at better than chance level whether they had received an active or sham tACS (McNemar’s test, *χ*^2^=1.125, p=.289). Stimulation was well tolerated, with only mild and transient side effects reported (e.g. itching and pain sensation close to the electrode, and phosphene perception).

**Table 1.** Contingency table of conditioning and reporting stimulation group.

|  | Report active | Report sham |
| --- | --- | --- |
| Condition active | 7 | 2 |
| Condition sham | 6 | 3 |

### EEG data acquisition

We recorded EEG signals during two Posner cueing task sessions. The EEG recording system included: two EEG amplifiers (g.USBamp), two driver (interface) boxes for active electrodes (g.GAMMAbox), an elastic EEG cap (g.EEGcap), and 32 active electrodes (g.ACTIVEelectrode, Ag/AgCl), one ground electrode (g.ACTIVEground, Ag/AgCl), and one reference ear clip (g.GAMMAearclip, Ag/AgCl). All components were manufactured by g.tec Medical Engineering GmbH, Schiedlberg, Austria. Conductive gel (g.GAMMAgel, g.tec Medical Engineering GmbH, Schiedlberg, Austria) was applied to the electrodes. Electrodes were mounted according to the international 10-20 system (Figure 1D) with A1 on the left ear as a common reference and Fpz on the forehead as a common ground. We used a microcontroller board (Arduino Uno, arduino.cc) to send a trigger signal from the computer for Posner cueing task to the EEG amplifier. EEG data were digitized at a sampling rate of 1200 Hz with a band-pass filter between 0.1 and 100 Hz and a notch filter at 50 Hz (48-52 Hz, to remove power line interferences). We use MATLAB Simulink to acquire EEG data from EEG amplifiers (Documentation (2020)).

### Data processing

#### Behavior data

Data processing for behavior data was conducted using Python, utilizing libraries including NumPy (Harris et al. (2020)) and pandas (McKinney (2010)). Analysis of reaction times was performed on correct responses. Early responses that were too fast to reflect genuine reactions (*<*100 ms after target onset), as well as extremely slow reaction times (*>*1 s after target onset) were removed. These rejections resulted in a loss of 5.3% ± 1.1% (M ± SD) of trials per subject. We categorized all trials into four types based on the cue-target combinations: endogenous cue followed by a valid target (Endo-Valid), endogenous cue followed by an invalid target (Endo-Invalid), exogenous cue followed by a valid target (Exo-Valid), and exogenous cue followed by an invalid target (Exo-Invalid).

#### EEG Preprocessing

The preprocessing and extraction of EEG signals were performed using Python, including MNE-Python (Gramfort et al. (2013)), NumPy (Harris et al. (2020)), pandas (McKinney (2010)), and SciPy (Virtanen et al. (2020)) libraries. The continuous EEG data were epoched by trials, from the onset of fixation to the end of the response stage. EEG epochs holding no responses or incorrect responses rejected by behavior data were removed. To clear the EEG data from artifacts like muscle, heart, eye blinks and eye movements, the remaining trials were fed into an independent component analysis (ICA) using a Picard method (Ablin et al. (2018)). We decomposed 32 independent components and manually rejected 6.2 ± 1.4 components (M ± SD) by screening topography, epochs image, event-related potential, power spectrum, and epoch variance of those components. These components were rejected due to muscle activity, eye blinks, heartbeats, and other noise artifacts that were identified in the screening. Finally, the data were re-referenced to the common average reference.

#### Event-related potentials (ERPs)

For ERP analyses, the epoched EEG data were low-pass filtered at 30 Hz (Zhang et al. (2023)) to attenuate high-frequency components and then linearly detrended. ERP epochs were extracted from − 0.1 to 0.5 s relative to target onset and baseline-corrected using the − 0.1 to 0 s interval. Based on prior work localizing visuospatial attention-related ERP components over parietal and occipital regions (Hillyard and Anllo-Vento (1998); Di Russo et al. (2003); Eimer (1999)), we focused on parieto-occipital electrodes (Pz, P5, P6, P7, P8, PO3, PO4, O1, O2), which are commonly used for ERP quantification in visuospatial attention tasks (Thut et al. (2006); Zani et al. (2020); Sadeghi and Nazari (2015); Szewczyk et al. (2022)). For each participant, single-trial epochs were averaged to obtain condition-specific ERPs, and signals were then averaged across the selected electrodes to yield one ROI-averaged ERP waveform per condition.

Because both ERP amplitude and latency were central outcomes, we quantified ERPs using predefined component time windows: 90–200 ms for N1 and 250–400 ms for P3 (Di Russo et al. (2003); Thut et al. (2006); Zani et al. (2020)). Component amplitude was quantified as the mean voltage within each time window, selected over peak-based measures for stability (Sur and Sinha (2009)). Component latency was quantified using a fractional-area latency approach (Sur and Sinha (2009); Di Russo et al. (2003)). Within each component window, we computed the rectified component area and defined latency as the time point at which the cumulative area reached 50% of the total area (FAL50), using linear interpolation. Where relevant, sensitivity checks using additional fractions (e.g., 25% and 75%) were performed to assess robustness (Sur and Sinha (2009)).

In addition to component-based analyses, we performed an exploratory time-resolved analysis to visualize when pre/post differences emerged over the ERP waveform. For each group and trial type, paired *t*-tests compared pre-vs. post-tACS ERP amplitudes at each time point, and the false discovery rate (FDR) was used to control for multiple comparisons across time (Benjamini and Hochberg (1995)).

#### Oscillatory power computation

For the computation of the oscillatory power of neuronal oscillations, we initially applied a linear detrend procedure to the epoched preprocessed EEG data. All channels were used, as changes in these indexes can occur across large regions of the brain (Fingelkurts and Fingelkurts (2014); Morales and Bowers (2022); Nakao et al. (2019)). EEG data were epoched to the cue-target interval (CTI) stage for each trial. Due to two lengths of CTI (*T*_*CTI*_ =0.5 or 1 s), the length of epoched data was also 0 to 0.5 s, or 0 to 1 s. We analyzed two frequency bands: alpha (8–12 Hz) and gamma (30–45 Hz). The mean of EEG data was subtracted from each segment before computing its power spectral density (PSD). We employed a multitaper method in the MNE-Python package to compute the PSD with DPSS tapers (Slepian (1978); Babadi and Brown (2014)). The frequency bandwidth was 8*/T*_*CTI*_. Only tapers with more than 90% spectral concentration within bandwidth were used. Oscillatory powers were computed by integrating PSD values over respected bands.

#### Detrended fluctuation analysis (DFA)

To quantify the long-range temporal correlations (LRTC) of EEG signals, we employed DFA to compute the *α*-value of a segment of EEG signal. We adhered to the methodology described by Peng et al. (1994), utilizing the fathon Python library (Bianchi (2020)). DFA was applied to EEG signals obtained during the CTI following endogenous cues, and the CTI following exogenous cues. The length of epochs were dependent on the length of CTI as *T*_*CTI*_ = 0.5 or 1 s, respectively. The EEG signal, as a discrete time series, was denoted as *x*(*i*), 1 ⩽ *i* ⩽ *N*. The initial step involved subtracting the mean from the time series and integrating it:

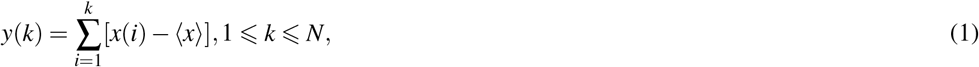

where ⟨*x*⟩ represents the average over the range [1, *N*]. Next, we divide *y*(*k*) into contiguous segments with length *n* (25 ⩽ *n* ⩽ 200). For each segment, the local linear trend *y*_*n*_(*k*) is determined by least-squares fit. The integrated time series *y*(*k*) is detrended by subtracting *y*_*n*_(*k*). The root-mean-square fluctuation is then computed by:

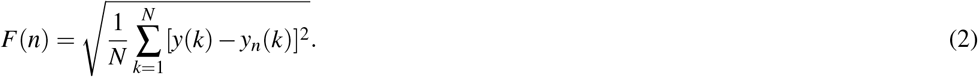

The computation of *F*(*n*) is repeated for a range of different scales *n*. The logarithm of *F*(*n*) against the logarithm of *n* is plotted, and the slope of the linear region, denoted as *α*, represents the scaling exponent:

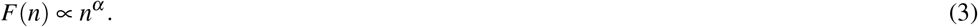

### Statistical analysis

Statistical inferences were conducted using Python, primarily utilizing SciPy (Virtanen et al. (2020)) and statsmodels (Seabold and Perktold (2010)).

#### Behavior data

Normality of reaction time (RT) distributions was assessed using the Shapiro–Wilk test (Shapiro and Wilk (1965)). The overall RT dataset and each trial-type subset yielded p*>*0.05; therefore, the null hypothesis of normality was not rejected. Our primary inferential model for RT was a 2×2×4 three-way repeated measures ANOVA with Group (sham vs. active) as a between-subject factor and Task time (pre vs. post) and Trial type (Endo-Valid, Endo-Invalid, Exo-Valid, Exo-Invalid) as within-subject factors (Mashal and Metzuyanim-Gorelick (2019); Du et al. (2022); Imbert et al. (2022)). The primary effect of interest was the offline stimulation effect, operationalized as the Group × Task time interaction and its modulation by Trial type (Group × Task time × Trial type).

Follow-up analyses were conducted to decompose significant omnibus interactions. Specifically, when the omnibus model indicated a trial-type-dependent offline effect, we performed planned contrasts within each trial type comparing pre–post changes between groups (ΔRT = post – pre; active vs. sham), with multiplicity across the four trial types controlled using Holm correction. Exploratory within-group pre vs. post comparisons were additionally evaluated using paired *t*-tests with Bonferroni correction (Armstrong (2014)).

#### Event-related potentials

To test stimulation-related effects on ERP component *amplitude* and *latency*, we conducted separate three-way repeated measures ANOVAs for N1 mean amplitude, P3 mean amplitude, N1 latency, and P3 latency. Each ANOVA included two within-subject factors (task time and trial type) and one between-subject factor (group) (Mashal and Metzuyanim-Gorelick (2019); Du et al. (2022); Imbert et al. (2022)). Post-hoc pairwise comparisons were restricted to planned contrasts with multiplicity control applied across the four trial types using the Holm correction. Additionally, we performed an exploratory time-resolved analysis to visualize when pre/post differences emerged over the ERP waveform. Paired *t*-tests compared pre-vs. post-tACS ERP amplitudes at each time point, with the false discovery rate (FDR) used to control for multiple comparisons across time (Benjamini and Hochberg (1995)).

#### Oscillatory power and LRTC

To measure the effects of tACS on both oscillatory power and LRTC (*α*-values) while controlling for multiple comparisons across the scalp, we utilized non-parametric cluster-based permutation analyses via MNE-Python (Davis et al. (2023); Murphy et al. (2020)). Clusters were defined as two or more adjacent electrodes at a p*<*0.05 in t-tests with identical signs of t-values. We used a Monte Carlo method with 2000 iterations to calculate the two-tailed significance probability. For significant electrode clusters, post hoc t-tests with a Bonferroni correction (Armstrong (2014)) were carried out to investigate pairwise differences between pre-/post-tACS and sham/active groups.

#### Sensitivity analysis

Because the sample size was fixed (*n* = 9 per group), we conducted an effect-size sensitivity analysis to contextualize the smallest effects our design could reliably detect (Giner-Sorolla et al. (2024)). We focused on the primary offline stimulation effect (Group × Time), operationalized as a between-group comparison of pre–post change scores (Δ = post – pre) with two-sided *α* = 0.05 and 80% power. Under these settings, the minimum detectable effect corresponds to a large standardized difference between groups (Cohen’s *d* ≈ 1.41). For trial-type–specific follow-up contrasts with multiplicity control across the four trial types (conservatively *α* ≈ 0.0125), the minimum detectable effect is larger (*d* ≈ 1.75). These values indicate that the study is powered to detect large effects, whereas smaller effects should be interpreted cautiously.

## RESULTS

### Reaction time is more reduced in post-active stimulation than post-sham stimulation

Reaction times for correct trials were first submitted to a 2×2×4 repeated-measures ANOVA, with factors of group, task time, and trial type. The omnibus analysis resulted in significant main effects of group (F(1,16)=19.26, p*<*.01,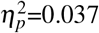), task time (F(1,16)=52.12, p*<*.01, 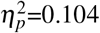), and trial type (F(3,48)=118.3, p*<*.01, 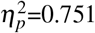). Moreover, there was a significant two-way interaction of group x task time (F(1,16)=24.72, p*<*.01, 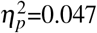). No significant main effect was observed in two-way interactions of group x trial type (F(3,48)=1.14, p=.34, 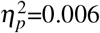), or task time x trial type (F(3,48)=2.35, p=.08, 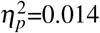). The three-way interaction of group x task time x trial type was significant (F(3,48)=3.12, p=.03, 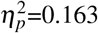). The three-way repeated-measures ANOVA represented that the reaction times were significantly influenced by the group, task time, and trial type, with a notable interaction between group and task time. This indicated that sham and active tACS intervention result in varied reaction time in post-tACS task compared to pre-tACS task. Due to the large effect size for the trial type, and the evidence that attention types (Dugué et al. (2020)) and cue validity (Dugué et al. (2017); Osaki et al. (2021)) had a significant impact on reaction time, we separated the analysis of each trial type and further studied the interaction between group and task times for each trial type (Figure 2).

**Figure 2.**
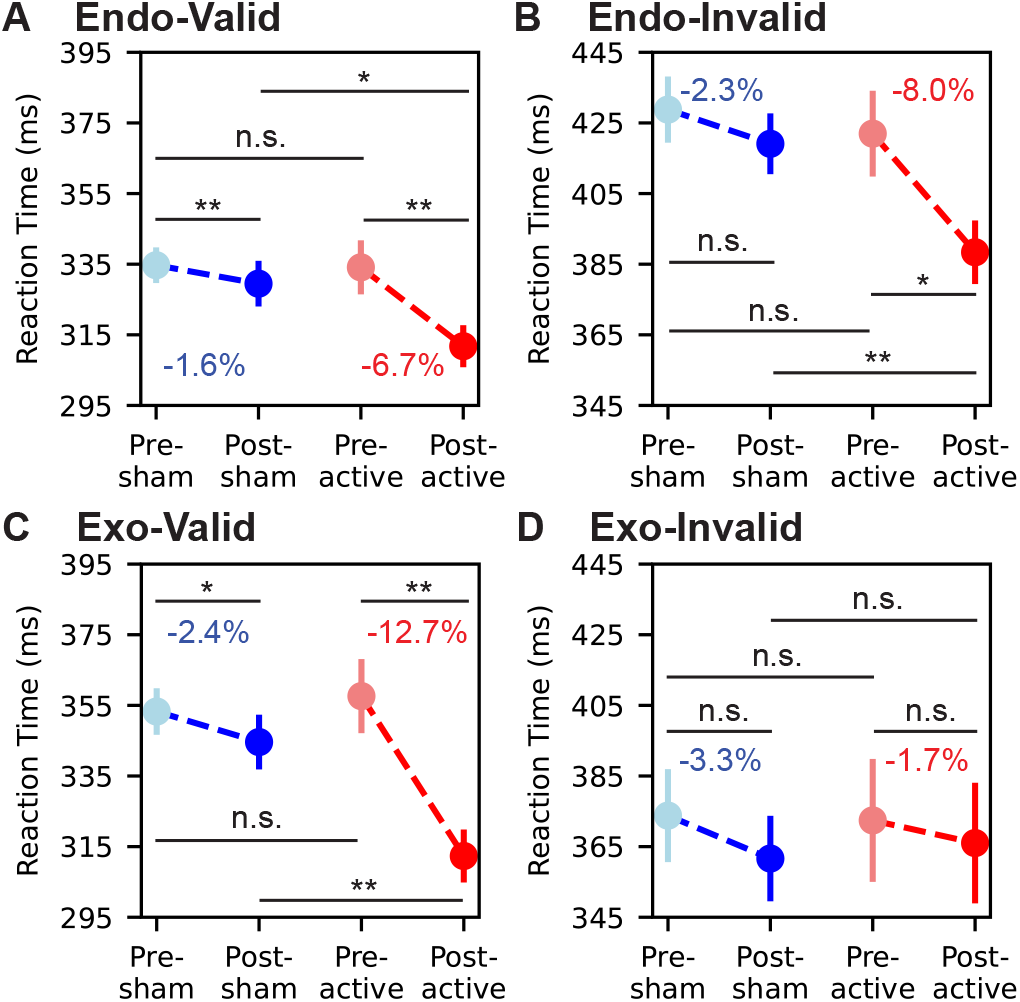
Reaction time change from pre-tACS task to post-tACS task. Trials presenting an endogenous cue followed and a valid target, in which the target appears on the same direction as the cue, is noted as ‘Endo-Valid’ (A). Same abbreviation rule for ‘Endo-Invalid’ (B), ‘Exo-Valid’ (C), and ‘Exo-Invalid’ (D). Light blue dots, blue dots, light red dots, and red dots denote the mean reaction time of the pre-tACS task of the sham group, the post-tACS task of the sham group, the pre-tACS task of the active group, and the post-tACS task of the active group, respectively. Henceforth we will use the abbreviation for groups and sessions as pre-sham, post-sham, pre-active, and post-active. The vertical lines passing through the dots denote 95%-confidence intervals of the mean reaction time. The percentage number in blue or red colors denotes the pre-post change of the mean reaction time. Four pairwise comparisons are computed to measure the statistical significance, which are: pre-sham and post-sham; pre-active and post-active; pre-sham and pre-active; post-sham and post-active. Pairwise comparisons used t-tests with a Bonferroni correction (*p*<*0.05, **p*<*0.01, n.s., not significant).

For the 2×2 repeated-measures ANOVA of Endo-Valid trials, main effects of group (F(1,16)=9.07, p*<*.01, 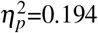) and task time (F(1,16)=16.98, p*<*.01, 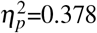) were significant. The main effect of two-way interaction of group x task time was also significant (F(1,16)=4.78, p=.04, 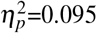). To further examine the effects of sham and active tACS on reaction time, we conducted a pairwise comparison among the reaction time in pre-sham, post-sham, pre-active and post-active tasks. Figure 2 summarizes pre/post changes by trial type. Pairwise t-tests confirmed the reaction time reduction in the sham group was significant (p=.02), from 334.7 ± 51.4 ms (M SD) in the pre-sham task to 329.5 ± 67.2 ms in the post-sham task. The reaction time reduction in the active group was also significant (p*<*.01), from 334.1 ± 80.3 ms in the pre-active task to 311.2 ± 60.8 ms in the post-active task. Moreover, the reaction time in the post-active task was significantly lower than in the post-sham task (p=.03).

For the 2×2 repeated-measures ANOVA of Endo-Invalid trials, main effects of group (F(1,16)=6.88, p=.02, 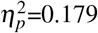) and task time (F(1,16)=11.72, p*<*.01, 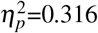) were significant. The main effect of two-way interaction of group x task time was also significant (F(1,16)=5.22, p*<*.03,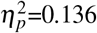). Pairwise t-tests (Figure 2B) confirmed the reaction time reduction in the sham group was not significant (p*>*.05), from 428.8 ± 94.1 ms in the pre-sham task to 419.1 ± 86.0 ms in the post-sham task. However, the reaction time reduction in the active group was significant (p=.01), from 422.0 ± 125.8 ms in the pre-active task to 388.4 ± 86.6 ms in the post-active task. Moreover, the reaction time in the post-active task was significantly lower than in the post-sham task (p*<*.01).

For the 2×2 repeated-measures ANOVA of Exo-Valid trials, main effects of group (F(1,16)=12.57, p*<*.01,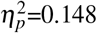) and task time (F(1,16)=39.51, p*<*.01, 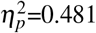) were significant. The main effect of two-way interaction of group x task time was also significant (F(1,16)=16.75, p*<*.01,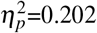). Pairwise t-tests (Figure 2C) confirmed the reaction time reduction in the sham group was significant (p=.01), from 353.3 ± 67.0 ms in the pre-sham task to 343.9 ± 80.7 ms in the post-sham task. The reaction time reduction in the active group was also significant (p*<*.01), from 357.6 ± 109.3 ms in the pre-active task to 312.3 ± 75.7 ms in the post-active task. Moreover, the reaction time in the post-active task was significantly lower than in the post-sham task (p*<*.01).

For the 2×2 repeated-measures ANOVA of Exo-Invalid trials, the main effects of neither group (F(1,16)=0.03, p=.86,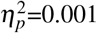) nor task time (F(1,16)=1.86, p=.19,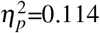) were significant. The main effect of two-way interaction of group x task time was also not significant (F(1,16)=0.49, p=.49,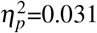). Although the mean reaction time reduced from 373.8 ± 68.8 ms to 361.6 ± 63.5 ms in the sham group and from 372.2 ± 90.5 to 366.0 ± 88.4 ms in the active group, pairwise t-tests (Figure 2D) revealed no significant difference among groups and task times (all p*>*.05). These results suggested that the after-effect of our stimulation protocol on reaction time was not uniform across trial types. Task performance in Endo-Valid, Endo-Invalid, and Exo-Valid trials represented significant enhancement after the active intervention of gamma tACS but not in Exo-Invalid trials.

### Diverse after-effects of active tACS on event-related potentials across trial types

We quantified the amplitude of ERP component as the mean voltage within an a priori N1 window (90–200 ms) and P3 window (250–400 ms), averaged across the parieto-occipital ROI electrodes. The latency of ERP component was quantified using the predefined latency metric described in the ERP Methods (computed within the same N1 and P3 windows). Amplitude and latency were analyzed in separate three-way repeated-measures ANOVA with *Group* (sham vs. active) as a between-subject factor and *Task time* (pre vs. post) and *Trial type* (Endo-Valid, Endo-Invalid, Exo-Valid, Exo-Invalid) as within-subject factors.

The omnibus ANOVA of N1 amplitude showed significant main effects of group (F(1,16)=6.71, p=.02,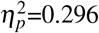), task time (F(1,16)=21.14, p*<*.01,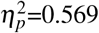), and trial type (F(3,48)=39.04, p*<*.01,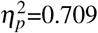). Moreover, there was a significant two-way interaction of group x task time (F(1,16)=22.74, p*<*.01,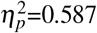). No significant main effect was observed in two-way interactions of group x trial type (F(3,48)=1.41, p=.25,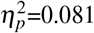), or task time x trial type (F(3,48)=1.12, p=.35,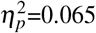). However, the three-way interaction of group x task time x trial type was significant (F(3,48)=3.04, p=.04,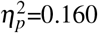). Planned follow-up contrasts (Holm-corrected across the four trial types) indicated that the offline effect on N1 amplitude was driven by the Endo-Valid condition: N1 amplitude changed from pre- to post-active in the active group relative to sham (p=.03), whereas no reliable N1 amplitude change was observed for Endo-Invalid, Exo-Valid, or Exo-Invalid trials. This pattern is summarized in the amplitude panel of Figure 3A.

**Figure 3.**
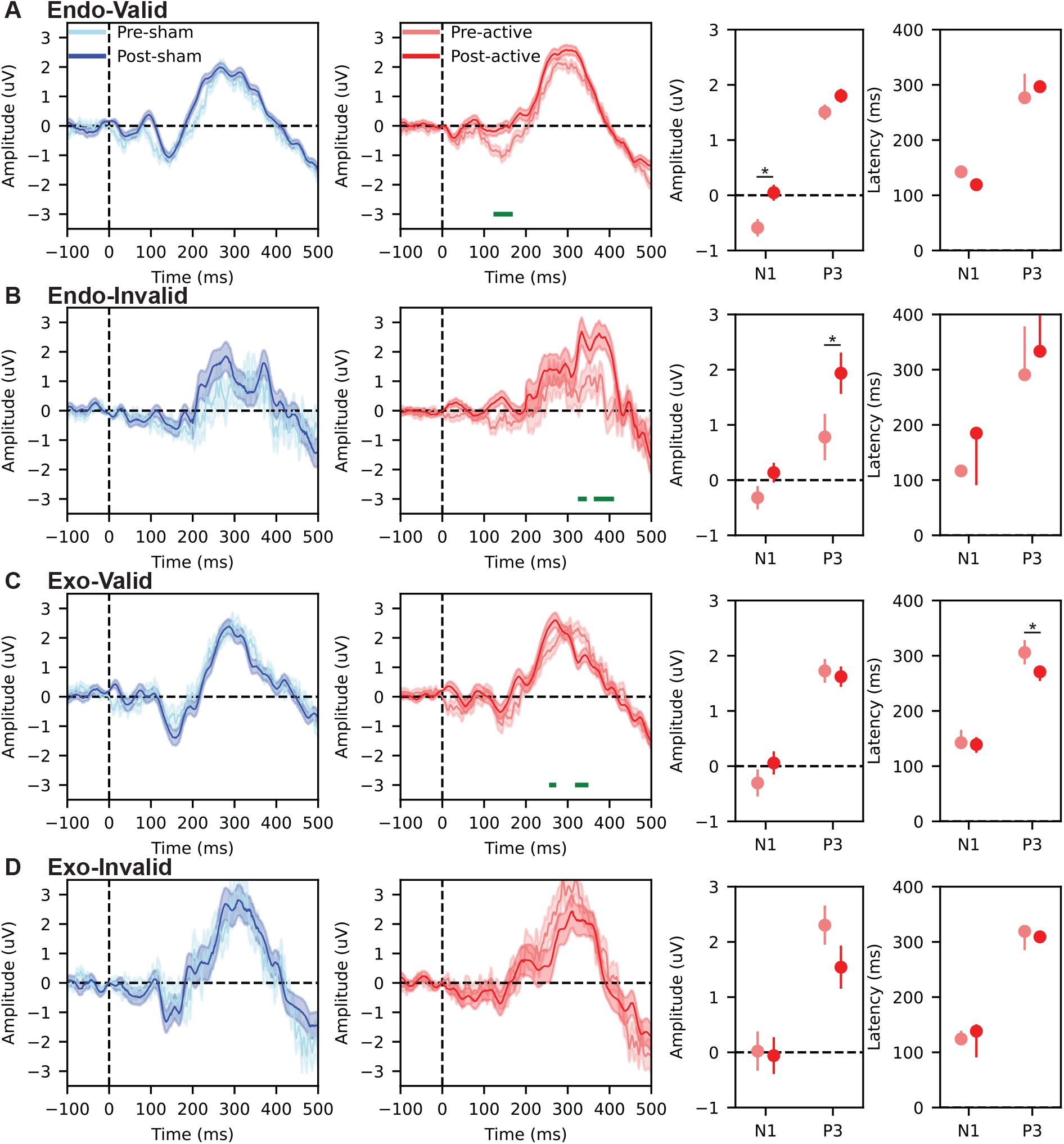
Event-related potentials (ERPs) time-locked to target onset and corresponding component-based amplitude and latency summaries. Panels (A)–(D) correspond to the four trial types (Endo-Valid, Endo-Invalid, Exo-Valid, Exo-Invalid). Within each row, the first two subplots show ROI-averaged ERP waveforms (parieto-occipital electrodes: Pz, P5, P6, P7, P8, PO3, PO4, O1, O2): sham group (pre vs. post) on the left and active group (pre vs. post) on the right. Solid lines represent group means and shaded areas indicate ± 1 SEM. Time 0 ms denotes target onset. Green horizontal bars indicate time points where pre vs. post waveforms differed in the exploratory time-resolved analysis (paired *t*-tests across time, FDR-corrected; Benjamini and Hochberg (1995)). The third subplot in each row summarizes component amplitude for N1 (90–200 ms) and P3 (250–400 ms) for the active group (pre vs. post), and the fourth subplot summarizes the corresponding component latency for N1 and P3 (active group: pre vs. post). Points and error bars show mean and 95%-confidence intervals of the mean. (*p*<*0.05)

For the ANOVA of ERP in the time window of P3 component, the omnibus analysis resulted in significant main effects of group (F(1,16)=19.38, p*<*.01,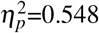), task time (F(1,16)=58.62, p*<*.01,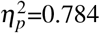), and trial type (F(3,48)=38.90, p*<*.01,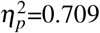). Moreover, there were significant two-way interaction of group x task time (F(1,16)=22.85, p*<*.01, 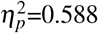), and task time x trial type (F(3,48)=3.22, p=.03, 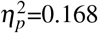). No significant main effect was observed in two-way interactions of group x trial type (F(3,48)=0.97, p=.41, 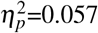). However, the three-way interaction of group x task time x trial type was significant (F(3,48)=3.14, p=.03, 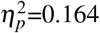). Planned follow-up contrasts (Holm-corrected) indicated that the offline effect on P3 mean amplitude was primarily expressed in the Endo-Invalid condition: P3 mean amplitude changed from pre- to post-active in the active group relative to sham (p=.02), whereas P3 mean-amplitude changes were not reliable in the remaining trial types. This pattern is summarized in the amplitude panel of Figure 3B.

Latency ANOVAs were performed separately for N1 and P3 using the same factor structure. For N1 latency, the omnibus effects and interactions were (group: F(1,16)=5.22, p=.04, 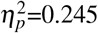; time: F(1,16)=6.2, p=.02, 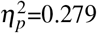; trial type: F(3,48)=8.69, p*<*.01, 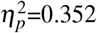; group x task time: F(1,16)=4.89, p=.04, 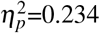; task time x trial type: F(3,48)=3.059, p=.04, 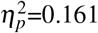; group x task time x trial type: F(3,48)=2.91, p=.04, 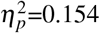), and planned contrasts did not indicate a reliable offline latency modulation across trial types after multiplicity control. For P3 latency, the omnibus ANOVA showed (group: F(1,16)=12.7, p*<*.01,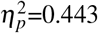; task time: F(1,16)=8.21, p=.01, 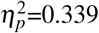; trial type: F(3,48)=5.84, p=0.002, 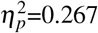; group x task time: F(1,16)=5.58, p=.03, 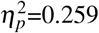; task time x trial type: F(3,48)=3.01, p=.04, 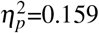; group x task time x trial type: F(3,48)=3.44, p=.02, 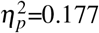), and planned contrasts (Holm-corrected) indicated that the offline latency effect was expressed in the Exo-Valid condition: P3 latency shortened from pre-to post-active in the active group relative to sham (p=.03), whereas P3 latency changes were not reliable in the other trial types. This pattern is summarized in the latency panel of Figure 3C.

In addition to the component-based measures above, we performed an exploratory time-resolved comparison of pre vs. post ERP amplitudes at each time point (paired *t*-tests, FDR-corrected across time). This analysis is visualized as green bars in Figure 3 and is intended to indicate *when* waveform differences occurred, rather than to define component latency. Consistent with the component-based results, time-resolved differences were observed in the active group for Endo-Valid trials within the N1 time range (Figure 3A), for Endo-Invalid trials within the P3 time range (Figure 3B), and for Exo-Valid trials within the P3 time range (Figure 3C), while no reliable time-resolved differences were observed for Exo-Invalid trials (Figure 3D).

Overall, the ERP results indicate trial-type-specific effects of gamma tACS, with dissociable modulation of component amplitude (Endo-Valid N1 and Endo-Invalid P3) and component latency (Exo-Valid P3), broadly paralleling the condition-dependent behavioral effects reported for reaction time.

### Decreasing of oscillatory power in alpha band but increasing in gamma band during CTI after active-tACS intervention

Further, we investigated the after-effects of gamma tACS on the oscillatory power during CTI period. Due to different roles of alpha and gamma oscillations as well as different mechanism of endogenous and exogenous attention (Riddle et al. (2019)), we separate the analysis of oscillatory power as frequency band x trial type, result in four combinations. For the oscillatory power in the alpha band in CTI following endogenous cue, non-parametric cluster-based permutation analysis revealed a significant decrease over the right hemisphere (pre-active vs. post-active, p=.021), while no significant difference was observed in comparisons of neither pre-sham vs. pre-active nor pre-sham vs. post-sham (Figure 4A, topographical maps). We further measured the average oscillatory power of electrodes in the significant cluster. As shown in the bar plot of Figure 4A, we found the alpha power in the significant cluster was significantly decreased after active-tACS (p=.03) and the alpha power was significantly lower in the post-active task than post-sham task (p*<*.01) (pre-sham vs. post-sham, pre-sham vs. pre-active, all p*>*.05).

**Figure 4.**
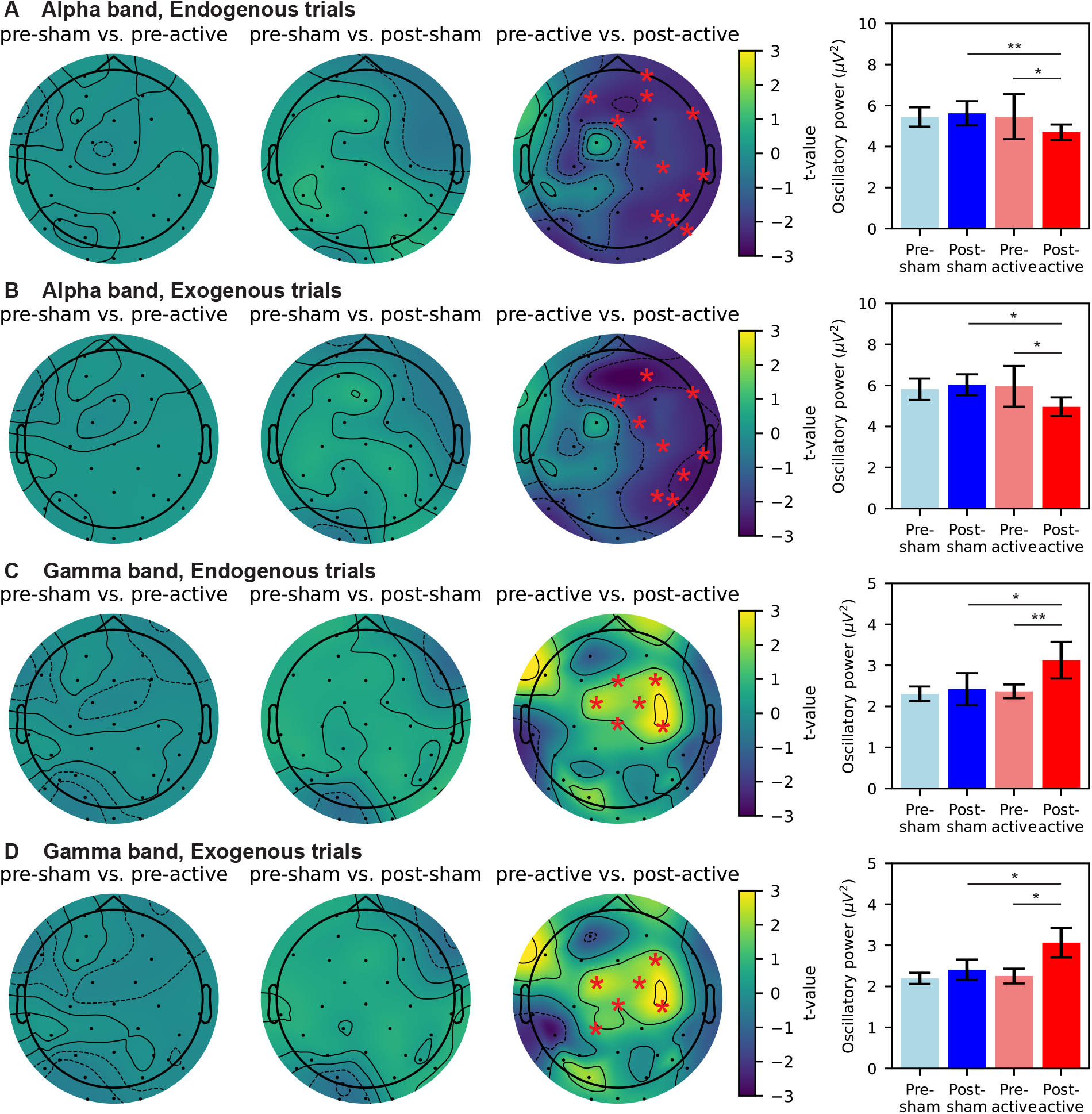
Change of oscillatory power during CTI due to sham/active tACS intervention. Panels from top to bottom represent oscillatory power in the alpha band of endogenous trials (A), alpha band of exogenous trials (B), gamma band of endogenous trials (C), and gamma band of exogenous trials (D), respectively. Topographical maps in each panel from left to right display differences in pre-sham vs. pre-active, pre-sham vs. post-sham, and pre-active vs. post-active, respectively. EEG electrodes forming significant clusters are marked by red stars (p*<*.05). Bar plots reflect the mean oscillatory power of electrodes marked in the corresponding topographical maps. Error bars denote 95%-confidence intervals of the mean oscillatory power. Pairwise comparisons in bar plots use t-tests with a Bonferroni correction (*p*<*0.05, **p*<*0.01).

For the oscillatory power in the alpha band in CTI following exogenous cue, non-parametric cluster-based permutation analysis revealed a significant decrease over the right hemisphere (pre-active vs. post-active, p=.017), while no significant difference was observed in comparisons of neither pre-sham vs. pre-active nor pre-sham vs. post-sham (Figure 4B, topographical maps). We further measured the average oscillatory power of electrodes in the significant cluster. As shown in the bar plot of Figure 4B, we found the alpha power in the significant cluster was significantly decreased after active-tACS (p=.04) and the alpha power was significantly lower in the post-active task than post-sham task (p=.02) (pre-sham vs. post-sham, pre-sham vs. pre-active, all p*>*.05).

For the oscillatory power in the gamma band in CTI following endogenous cue, non-parametric cluster-based permutation analysis revealed a significant increase over the frontal and central sulcus regions (pre-active vs. post-active, p=.004), while no significant difference was observed in comparisons of neither pre-sham vs. pre-active nor pre-sham vs. post-sham (Figure 4C, topographical maps). We further measured the average oscillatory power of electrodes in the significant cluster. As shown in the bar plot of Figure 4C, we found the gamma power in the significant cluster was significantly increased after active-tACS (p=.01) and the gamma power was significantly higher in the post-active task than post-sham task (p=.01) (pre-sham vs. post-sham, pre-sham vs. pre-active, all p*>*.05).

For the oscillatory power in the gamma band in CTI following exogenous cue, non-parametric cluster-based permutation analysis revealed a significant increase over the frontal and central sulcus regions (pre-active vs. post-active, p=.009), while no significant difference was observed in comparisons of neither pre-sham vs. pre-active nor pre-sham vs. post-sham (Figure 4D, topographical maps). We further measured the average oscillatory power of electrodes in the significant cluster. As shown in the bar plot of 4D, we found the gamma power in the significant cluster was significantly increased after active-tACS (p=.02) and the gamma power was significantly higher in the post-active task than post-sham task (p=.04) (pre-sham vs. post-sham, pre-sham vs. pre-active, all p*>*.05). Our findings revealed significant after-effects of gamma tACS on alpha band and gamma band oscillatory power. Active gamma tACS suppressed alpha power on the ipsilateral side of tACS intervention while increasing gamma power over frontal and central sulcus regions. Moreover, such modulation was consistent across endogenous and exogenous trials.

### Decreasing in long-range temporal correlations following active-tACS

We applied detrended fluctuation analysis (DFA) (Peng et al. (1994)) to compute the scaling exponent *α*-value of EEG signal that could assess LRTC changes induced by gamma tACS. The EEG data used for DFA was the segment of CTI stages following endogenous cue or exogenous cue, the same periods as the investigation of the oscillatory power.

For the long-range temporal correlations of EEG signal in the CTI following endogenous cue, non-parametric cluster-based permutation analysis revealed a significant decrease over the right hemisphere (pre-active vs. post-active, p=.043), while no significant difference was observed in comparisons of neither pre-sham vs. pre-active nor pre-sham vs. post-sham (Figure 5A, topographical maps). We further measured the average *α*-value of electrodes in the significant cluster. As shown in the bar plot of 5A, we found the *α*-value in the significant cluster was significantly decreased after active-tACS (p=.01) and the *α*-value was significantly lower in the post-active task than post-sham task (p=.02) (pre-sham vs. post-sham, pre-sham vs. pre-active, all p*>*.05).

**Figure 5.**
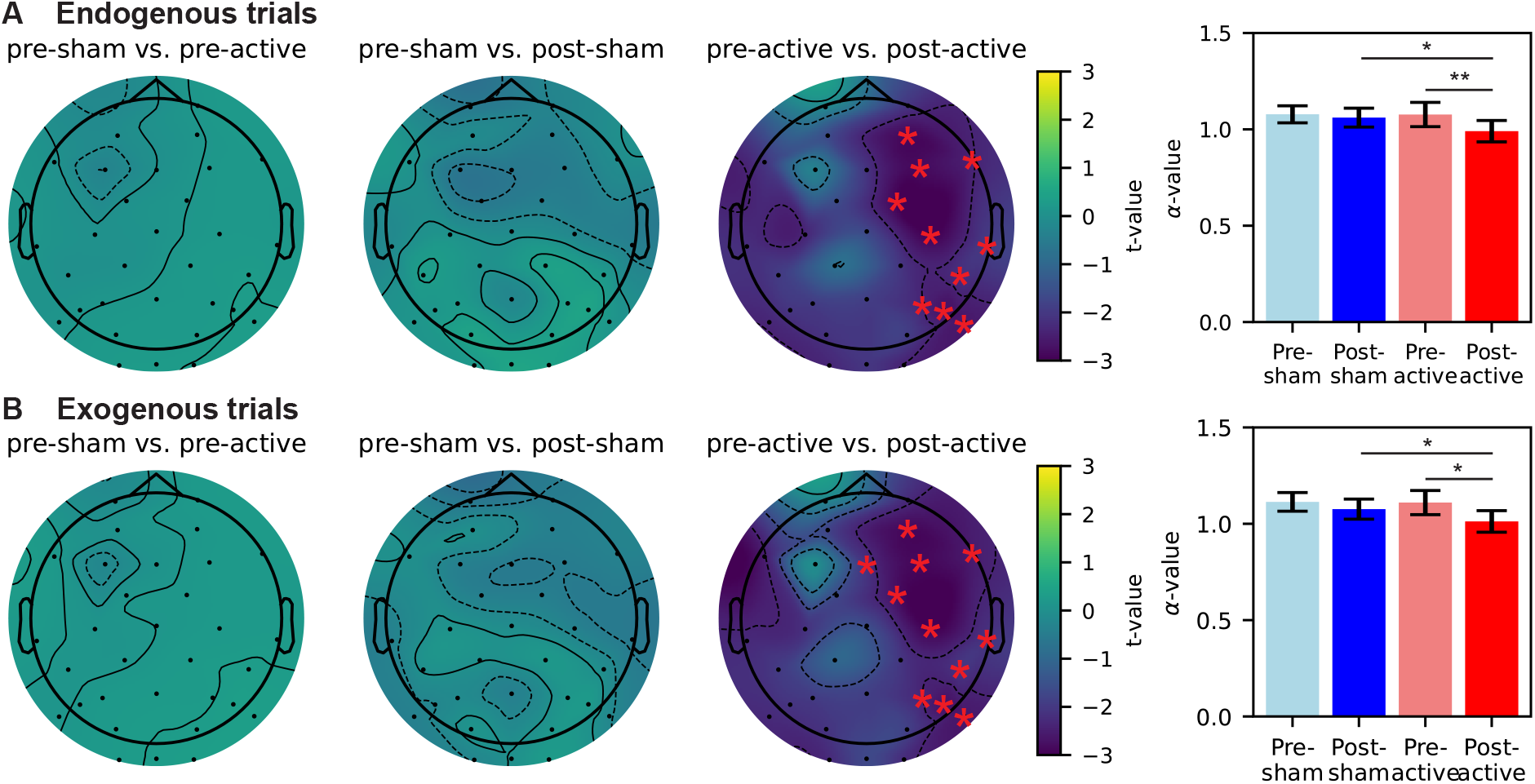
Change of long-range temporal correlations during CTI due to sham/active tACS intervention. (A) Change of *α*-value in the endogenous trials. (B) Change of *α*-value in the exogenous trials. Topographical maps in each panel from left to right display differences in pre-sham vs. pre-active, pre-sham vs. post-sham, and pre-active vs. post-active, respectively. EEG electrodes forming significant clusters are marked by red stars (p*<*.05). Bar plots reflect the mean *α*-value of electrodes marked in the corresponding topographical maps. Error bars denote 95%-confidence intervals of the mean *α*-value. Pairwise comparisons in bar plots use t-tests with a Bonferroni correction (*p*<*0.05, **p*<*0.01).

For the long-range temporal correlations of EEG signal in the CTI following exogenous cue, non-parametric cluster-based permutation analysis revealed a significant decrease over the right hemisphere (pre-active vs. post-active, p=.031), while no significant difference was observed in comparisons of neither pre-sham vs. pre-active nor pre-sham vs. post-sham (Figure 5B, topographical maps). We further measured the average *α*-value of electrodes in the significant cluster. As shown in the bar plot of Figure 5B, we found the *α*-value in the significant cluster was significantly decreased after active-tACS (p=.04) and the *α*-value was significantly lower in the post-active task than post-sham task (p=.02) (pre-sham vs. post-sham, pre-sham vs. pre-active, all p*>*.05). We revealed an after-effect of gamma tACS on LRTC during CTI. The *α*-value, which was an indicator of LRTC, was significantly decreased on the ipsilateral side of tACS intervention after active tACS, but not after sham tACS. Similar to our findings of oscillatory power, such modulation effect in CTI was consistent across endogenous and exogenous trials.

## DISCUSSION

The present study examined whether parieto-occipital 40 Hz tACS produces *offline* (post-stimulation) changes in visuo-spatial attention. At the behavioral level, active stimulation was associated with a reduction in reaction time (RT) from pre-to post-stimulation, with effects that were not uniform across cueing conditions (Figure 2). At the neural level, we observed post-stimulation modulation of target-evoked ERPs (N1/P3 amplitude and latency; Figure 3), accompanied by band-specific changes in cue–target interval (CTI) oscillatory power (alpha decrease and gamma increase; Figure 4) and reduced long-range temporal correlations (LRTC; Figure 5). Importantly, the ERP effects were expressed primarily in the same trial types that showed behavioral improvement, whereas the CTI power and LRTC changes were more consistent across endogenous and exogenous trials, suggesting a combination of condition-specific effects on target processing and more general changes in preparatory network dynamics.

### Behavioral after-effects and condition specificity

The sham group showed a modest RT reduction from the pre-to post-session, consistent with practice/familiarization effects commonly observed in reaction time tasks (Ghuntla et al. (2014); Clark et al. (2015)). Against this background, the active group exhibited a larger RT reduction following stimulation, indicating that 40 Hz tACS may contribute to offline performance changes beyond practice alone (Figure 2). The pattern was not uniform across conditions: RT reductions were evident in endogenous trials (valid and invalid) and in exogenous valid trials, whereas exogenous invalid trials did not show a reliable improvement. Prior *online* studies reported that gamma-frequency tACS can modulate cueing performance in a condition-dependent manner, including differential effects for endogenous versus exogenous attention (Hopfinger et al. (2017); Kasten et al. (2020)). Our results extend this line of work by suggesting that such modulation can persist after stimulation, while remaining selective across trial types.

One interpretation of this selectivity is that offline gamma tACS preferentially benefits processing regimes that are strongly engaged by the task structure (e.g., target processing under higher expectancy in valid trials) and/or by the control demands of endogenous orienting. However, given the modest sample size and the between-group design, this condition specificity should be treated as preliminary and in need of replication with confirmatory analyses. Mechanistically, offline effects may reflect stimulation-induced plastic changes rather than transient entrainment (Zaehle et al. (2010); Vossen et al. (2015)), and may also depend on endogenous network state (Schmidt et al. (2014); Krause et al. (2022)). Consistent with this broader view, stimulation has been shown to interact with event-related rhythmic activity in ways that are not fully captured by phase synchronization accounts (Wischnewski and Schutter (2017)).

### Linking behavioral changes to ERP, oscillatory power, and LRTC

The electrophysiological results provide convergent evidence that offline 40 Hz tACS can modulate both time-locked target processing and ongoing preparatory dynamics. First, ERP analyses indicated modulation of N1 and P3 components after active stimulation (Figure 3). N1 has been linked to early attentional selection and sensory processing of visual targets (Di Russo et al. (2003)), whereas P3 amplitude is often interpreted as reflecting attentional resource allocation and stimulus evaluation processes (Chueh et al. (2017); Chang et al. (2015)). Importantly, P3 latency is commonly used as an index of processing speed, with shorter latencies often associated with faster responses (Miniussi et al. (1999)). In our data, P3 latency (FAL50) was modulated after stimulation (Figure 3C), and the ERP effects were expressed in the same trial types that showed RT reduction. Together with prior work linking RT to ERP amplitude and latency measures (Krieger and Dillbeck (1987); Kreegipuu and Allik (2007)), this pattern supports the interpretation that offline gamma tACS may alter target-processing stages that contribute to behavioral response speed.

Second, CTI oscillatory power changes suggest that stimulation affected preparatory attentional state in a frequency-specific manner (Figure 4). We observed a post-stimulation decrease in alpha power on the stimulation-ipsilateral side across both endogenous and exogenous trials, alongside a post-stimulation increase in gamma power over broader regions. Alpha-band dynamics during attention have been linked to sensory suppression and selective inhibition (Foxe and Snyder (2011)), and combined alpha/gamma measures have been related to response speed in attentional tasks (Fahimi et al. (2018)). Cross-frequency interactions provide one plausible framework for why gamma-frequency stimulation may be accompanied by alpha-band changes (Sotero (2016)). Notably, these oscillatory changes were less condition-specific than the ERP effects, which may indicate that stimulation altered a more general preparatory mode (CTI state) while selectively impacting target processing (ERP) in the conditions that showed behavioral improvement.

Third, we observed a reduction in LRTC (lower DFA *α*-values) over the stimulation-ipsilateral hemisphere during CTI (Figure 5). DFA-based *α*-values are widely used to quantify scale-free temporal structure in time series (Peng et al. (1994); Gebber et al. (2006)). Prior work has linked stronger long-range temporal correlations in beta/gamma activity to poorer sustained visual attention (Irrmischer et al. (2018)). Consistent with this literature, the combined pattern of faster RT (in most conditions) and reduced LRTC suggests that offline gamma tACS may shift ongoing dynamics toward a less persistent, potentially more flexible regime during preparatory periods.

### Mechanistic considerations, limitations, and future directions

Although our findings are broadly consistent with the view that rhythmic stimulation can modulate oscillatory network activity beyond the stimulation period (Veniero et al. (2015)), several mechanistic and methodological issues warrant caution. First, the spatial specificity of conventional-intensity scalp stimulation remains debated, and stimulation effects may reflect a mixture of on-target cortical modulation, field spread, and peripheral co-stimulation (Vöröslakos et al. (2018)). Second, tACS effects are known to depend on endogenous oscillatory state and ongoing entrainment strength (Schmidt et al. (2014); Krause et al. (2022)), which could contribute to inter-individual variability and condition specificity. Third, the present study used a small sample size (n=9/group) in a between-group design while applying multiple analyses across behavioral conditions and EEG metrics. Consistent with the sensitivity analysis, the present design was primarily powered to detect large Group x Task time effects, and smaller effects remain uncertain. While we controlled specific comparisons (e.g., FDR in targeted tests), the combination of modest sample size and analytical breadth increases uncertainty in effect-size estimation and generalizability. Therefore, the present results should be interpreted as preliminary evidence motivating confirmatory replication.

Future work should test these effects in larger, preregistered studies with clearly specified primary outcomes (amplitude and latency endpoints for ERP components), more strict control of stimulation-related sensations and blinding, and complementary approaches to improve montage interpretability (e.g., electric-field modeling and/or individualized targeting). Within-subject or crossover designs may also help to reduce between-group variance and increase sensitivity to offline effects.

## CONCLUSIONS

In this single-blind, sham-controlled between-group study, parieto-occipital 40 Hz tACS was associated with offline reductions in reaction time in a visuo-spatial attention task, with effects that were condition-dependent. Convergent EEG results indicated post-stimulation modulation of target-evoked ERPs (N1/P3 amplitude and latency) alongside frequency-specific changes in CTI oscillatory power (alpha decrease and gamma increase) and reduced LRTC. Taken together, these findings provide preliminary evidence that offline gamma tACS can modulate both behavioral performance and electrophysiological signatures of attentional processing.

## DATA AVAILABILITY

The de-identified behavioral dataset and derived EEG measures supporting the findings of this study are publicly available on OpenNeuro (accession number: ds007043; DOI: 10.18112/openneuro.ds007043.v1.0.0). All shared materials have been de-identified to protect participant privacy.

## REFERENCES

Abellaneda-Pérez, K., Vaqué-Alcázar, L., Perellón-Alfonso, R., Bargalló, N., Kuo, M.-F., Pascual-Leone, A., Nitsche, M. A., and Bartrés-Faz, D. (2020). Differential tdcs and tacs effects on working memory-related neural activity and resting-state connectivity. Frontiers in Neuroscience, 13.

Ablin, P., Cardoso, J.-F., and Gramfort, A. (2018). Faster independent component analysis by preconditioning with hessian approximations. IEEE Transactions on Signal Processing, 66(15):4040–4049.

Antal, A., Alekseichuk, I., Bikson, M., Brockmöller, J., Brunoni, A., Chen, R., Cohen, L., Dowthwaite, G., Ellrich, J., Flöel, A., Fregni, F., George, M., Hamilton, R., Haueisen, J., Herrmann, C., Hummel, F., Lefaucheur, J., Liebetanz, D., Loo, C., McCaig, C., Miniussi, C., Miranda, P., Moliadze, V., Nitsche, M., Nowak, R., Padberg, F., Pascual-Leone, A., Poppendieck, W., Priori, A., Rossi, S., Rossini, P., Rothwell, J., Rueger, M., Ruffini, G., Schellhorn, K., Siebner, H., Ugawa, Y., Wexler, A., Ziemann, U., Hallett, M., and Paulus, W. (2017). Low intensity transcranial electric stimulation: Safety, ethical, legal regulatory and application guidelines. Clinical Neurophysiology, 128(9):1774–1809.

Armstrong, R. A. (2014). When to use the bonferroni correction. Ophthalmic and Physiological Optics, 34(5):502–508.

Babadi, B. and Brown, E. N. (2014). A review of multitaper spectral analysis. IEEE Transactions on Biomedical Engineering, 61(5):1555–1564.

Benjamini, Y. and Hochberg, Y. (1995). Controlling the false discovery rate: A practical and powerful approach to multiple testing. Journal of the Royal Statistical Society: Series B (Methodological), 57(1):289–300.

Bianchi, S. (2020). fathon: A python package for a fast computation of detrendend fluctuation analysis and related algorithms. Journal of Open Source Software, 5(45):1828.

Bisley, J. W. and Goldberg, M. E. (2010). Attention, intention, and priority in the parietal lobe. Annual Review of Neuroscience, 33(1):1–21. PMID: 20192813.

Bosman, C. A., Schoffelen, J.-M., Brunet, N., Oostenveld, R., Bastos, A. M., Womelsdorf, T., Rubehn, B., Stieglitz, T., Weerd, P. D., and Fries, P. (2012). Attentional stimulus selection through selective synchronization between monkey visual areas. Neuron, 75(5):875–888.

Braga, M., Barbiani, D., Emadi Andani, M., Villa-Sánchez, B., Tinazzi, M., and Fiorio, M. (2021). The role of expectation and beliefs on the effects of non-invasive brain stimulation. Brain Sciences, 11(11).

Chang, Y.-K., Pesce, C., Chiang, Y.-T., Kuo, C.-Y., and Fong, D.-Y. (2015). Antecedent acute cycling exercise affects attention control: an erp study using attention network test. Frontiers in Human Neuroscience, 9:156.

Chen, X., Shi, X., Wu, Y., Zhou, Z., Chen, S., Han, Y., and Shan, C. (2022). Gamma oscillations and application of 40-hz audiovisual stimulation to improve brain function. Brain and Behavior, 12(12):e2811.

Chueh, T.-Y., Huang, C.-J., Hsieh, S.-S., Chen, K.-F., Chang, Y.-K., and Hung, T.-M. (2017). Sports training enhances visuo-spatial cognition regardless of open-closed typology. PeerJ, 5:e3336.

Clark, K., Appelbaum, L. G., van den Berg, B., Mitroff, S. R., and Woldorff, M. G. (2015). Improvement in visual search with practice: Mapping learning-related changes in neurocognitive stages of processing. Journal of Neuroscience, 35(13):5351–5359.

Corbetta, M. and Shulman, G. L. (2002). Control of goal-directed and stimulus-driven attention in the brain. Nature Reviews Neuroscience, 3(3):201–215.

Davis, M.-C., Fitzgerald, P. B., Bailey, N. W., Sullivan, C., Stout, J. C., Hill, A. T., and Hoy, K. E. (2023). Effects of medial prefrontal transcranial alternating current stimulation on neural activity and connectivity in people with huntington’s disease and neurotypical controls. Brain Research, 1811:148379.

Di Russo, F., Martínez, A., and Hillyard, S. A. (2003). Source Analysis of Event-related Cortical Activity during Visuo-spatial Attention. Cerebral Cortex, 13(5):486–499.

Documentation, S. (2020). Simulation and model-based design.

Du, X. D., Li, Z., Yuan, N., Yin, M., Zhao, X. L., Lv, X. L., Zou, S. Y., Zhang, J., Zhang, G. Y., Li, C. W., Pan, H., Yang, L., Wu, S. Q., Yue, Y., Wu, Y. X., and Zhang, X. Y. (2022). Delayed improvements in visual memory task performance among chronic schizophrenia patients after high-frequency repetitive transcranial magnetic stimulation. World J Psychiatry, 12(9):1169–1182.

Dugué, L., Merriam, E. P., Heeger, D. J., and Carrasco, M. (2017). Specific Visual Subregions of TPJ Mediate Reorienting of Spatial Attention. Cerebral Cortex, 28(7):2375–2390.

Dugué, L., Merriam, E. P., Heeger, D. J., and Carrasco, M. (2020). Differential impact of endogenous and exogenous attention on activity in human visual cortex. Scientific Reports, 10(1):21274.

Eimer, M. (1999). Attending to quadrants and ring-shaped regions: Erp effects of visual attention in different spatial selection tasks. Psychophysiology, 36(4):491–503.

Elyamany, O., Leicht, G., Herrmann, C. S., and Mulert, C. (2021). Transcranial alternating current stimulation (tacs): from basic mechanisms towards first applications in psychiatry. European Archives of Psychiatry and Clinical Neuroscience, 271(1):135–156.

Fahimi, F., Goh, W. B., Lee, T.-S., and Guan, C. (2018). Neural indexes of attention extracted from eeg correlate with elderly reaction time in response to an attentional task. In Proceedings of the 3rd International Conference on Crowd Science and Engineering, ICCSE’18, New York, NY, USA. Association for Computing Machinery.

Fingelkurts, A. A. and Fingelkurts, A. A. (2014). Eeg oscillatory states: Universality, uniqueness and specificity across healthy-normal, altered and pathological brain conditions. PLOS ONE, 9(2):1–20.

Foxe, J. J. and Snyder, A. C. (2011). The role of alpha-band brain oscillations as a sensory suppression mechanism during selective attention. Frontiers in Psychology, 2.

Gebber, G. L., Orer, H. S., and Barman, S. M. (2006). Fractal noises and motions in time series of presympathetic and sympathetic neural activities. Journal of Neurophysiology, 95(2):1176–1184. PMID: 16306172.

Ghuntla, T. P., Mehta, H. B., Gokhale, P. A., and Shah, C. J. (2014). Influence of practice on visual reaction time. Journal of Mahatma Gandhi Institute of Medical Sciences, 19:119 – 122.

Giner-Sorolla, R., Montoya, A. K., Reifman, A., Carpenter, T., Lewis, Neil A., J., Aberson, C. L., Bostyn, D. H., Conrique, B. G., Ng, B. W., Schoemann, A. M., and Soderberg, C. (2024). Power to detect what? considerations for planning and evaluating sample size. Personality and Social Psychology Review, 28(3):276–301.

Gramfort, A., Luessi, M., Larson, E., Engemann, D. A., Strohmeier, D., Brodbeck, C., Goj, R., Jas, M., Brooks, T., Parkkonen, L., and Hämäläinen, M. S. (2013). MEG and EEG data analysis with MNE-Python. Frontiers in Neuroscience, 7(267):1–13.

Gruber, T., Müller, M. M., Keil, A., and Elbert, T. (1999). Selective visual-spatial attention alters induced gamma band responses in the human eeg. Clinical Neurophysiology, 110(12):2074–2085.

Han, C., Shapley, R., and Xing, D. (2022). Gamma rhythms in the visual cortex: functions and mechanisms. Cognitive Neurodynamics, 16:745–756.

Harris, C. R., Millman, K. J., van der Walt, S. J., Gommers, R., Virtanen, P., Cournapeau, D., Wieser, E., Taylor, J., Berg, S., Smith, N. J., Kern, R., Picus, M., Hoyer, S., van Kerkwijk, M. H., Brett, M., Haldane, A., del Río, J. F., Wiebe, M., Peterson, P., Gérard-Marchant, P., Sheppard, K., Reddy, T., Weckesser, W., Abbasi, H., Gohlke, C., and Oliphant, T. E. (2020). Array programming with NumPy. Nature, 585(7825):357–362.

Herwig, U., Satrapi, P., and Schönfeldt-Lecuona, C. (2003). Using the international 10-20 eeg system for positioning of transcranial magnetic stimulation. Brain Topography, 16(2):95–99.

Hillyard, S. A. and Anllo-Vento, L. (1998). Event-related brain potentials in the study of visual selective attention. Proceedings of the National Academy of Sciences, 95(3):781–787.

Homan, R. W., Herman, J., and Purdy, P. (1987). Cerebral location of international 10–20 system electrode placement. Electroencephalography and Clinical Neurophysiology, 66(4):376–382.

Hopfinger, J. B., Parsons, J., and Fröhlich, F. (2017). Differential effects of 10-hz and 40-hz transcranial alternating current stimulation (tacs) on endogenous versus exogenous attention. Cogn Neurosci, 8(2):102–111.

Huang, Y., Datta, A., Bikson, M., and Parra, L. (2019). Realistic volumetric-approach to simulate transcranial electric stimulation – roast – a fully automated open-source pipeline. Journal of Neural Engineering, 16(5).

Huang, Y., Parra, L., and Haufe, S. (2016). The new york head - a precise standardized volume conductor model for eeg source localization and tes targeting. NeuroImage, 140:150–162.

Huang, Y. and Datta, A. and Bikson, M. and Parra, L.C. (2018). ROAST: an open-source, fully-automated, Realistic vOlumetric-Approach-based Simulator for TES. Proceedings of the 40th Annual International Conference of the IEEE Engineering in Medicine and Biology Society, Honolulu, HI, July 2018.

Imbert, L., Moirand, R., Bediou, B., Koenig, O., Chesnoy, G., Fakra, E., and Brunelin, J. (2022). A single session of bifrontal tdcs can improve facial emotion recognition in major depressive disorder: An exploratory pilot study. Biomedicines, 10(10).

Irrmischer, M., Poil, S.-S., Mansvelder, H. D., Intra, F. S., and Linkenkaer-Hansen, K. (2018). Strong long-range temporal correlations of beta/gamma oscillations are associated with poor sustained visual attention performance. European Journal of Neuroscience, 48(8):2674–2683.

Jia, X., Smith, M. A., and Kohn, A. (2011). Stimulus selectivity and spatial coherence of gamma components of the local field potential. Journal of Neuroscience, 31(25):9390–9403.

Kasten, F. H., Wendeln, T., Stecher, H. I., and Herrmann, C. S. (2020). Hemisphere-specific, differential effects of lateralized, occipital–parietal α-versus γ-tacs on endogenous but not exogenous visual-spatial attention. Scientific Reports, 10(1):12270.

Klimesch, W. (2012). Alpha-band oscillations, attention, and controlled access to stored information. Trends in Cognitive Sciences, 16(12):606–617.

Krause, M. R., Vieira, P. G., Thivierge, J.-P., and Pack, C. C. (2022). Brain stimulation competes with ongoing oscillations for control of spike timing in the primate brain. PLOS Biology, 20(5):1–27.

Kreegipuu, K. and Allik, J. (2007). Detection of motion onset and offset: reaction time and visual evoked potential analysis. Psychological Research, 71:703–708.

Krieger, D. and Dillbeck, M. (1987). High frequency scalp potentials evoked by a reaction time task. Electroencephalography and Clinical Neurophysiology, 67(3):222–230.

Lorist, M. and Snel, J. (1998). Nicotine, Caffeine and Social Drinking: Behaviour and Brain Function. Routledge, 1st edition.

Magazzini, L. and Singh, K. D. (2018). Spatial attention modulates visual gamma oscillations across the human ventral stream. NeuroImage, 166:219–229.

Mashal, N. and Metzuyanim-Gorelick, S. (2019). New information on the effects of transcranial direct current stimulation on n-back task performance. Experimental Brain Research, 237(5):1315–1324.

McDermott, B., Porter, E., Hughes, D., McGinley, B., Lang, M., O’Halloran, M., and Jones, M. (2018). Gamma band neural stimulation in humans and the promise of a new modality to prevent and treat alzheimer’s disease. Journal of Alzheimer’s Disease, 65(2):363–392.

McKinney, W. (2010). Data structures for statistical computing in python. In Proceedings of the 9th Python in Science Conference, volume 445, pages 51–56. Austin, TX.

Meisel, C., Bailey, K., Achermann, P., and Plenz, D. (2017). Decline of long-range temporal correlations in the human brain during sustained wakefulness. Scientific Reports, 7(1):11825.

Miniussi, C., Wilding, E. L., Coull, J. T., and Nobre, A. C. (1999). Orienting attention in time: Modulation of brain potentials. Brain, 122(8):1507–1518.

Morales, S. and Bowers, M. E. (2022). Time-frequency analysis methods and their application in developmental eeg data. Developmental Cognitive Neuroscience, 54:101067.

Murphy, O., Hoy, K., Wong, D., Bailey, N., Fitzgerald, P., and Segrave, R. (2020). Transcranial random noise stimulation is more effective than transcranial direct current stimulation for enhancing working memory in healthy individuals: Behavioural and electrophysiological evidence. Brain Stimulation, 13(5):1370–1380.

Nakao, T., Miyagi, M., Hiramoto, R., Wolff, A., Gomez-Pilar, J., Miyatani, M., and Northoff, G. (2019). From neuronal to psychological noise – long-range temporal correlations in eeg intrinsic activity reduce noise in internally-guided decision making. NeuroImage, 201:116015.

Osaki, S., Amimoto, K., Miyazaki, Y., Tanabe, J., and Yoshihiro, N. (2021). Investigating the characteristics of covert unilateral spatial neglect using the modified posner task: A single-subject design study. Progress in Rehabilitation Medicine, 6.

Palva, J. M., Zhigalov, A., Hirvonen, J., Korhonen, O., Linkenkaer-Hansen, K., and Palva, S. (2013). Neuronal long-range temporal correlations and avalanche dynamics are correlated with behavioral scaling laws. Proceedings of the National Academy of Sciences, 110(9):3585–3590.

Park, S., Choi, D., Yi, J., Lee, S., Lee, J. E., Choi, B., Lee, S., and Kyung, G. (2017). Effects of display curvature, display zone, and task duration on legibility and visual fatigue during visual search task. Applied Ergonomics, 60:183–193.

Peng, C.-K., Buldyrev, S. V., Havlin, S., Simons, M., Stanley, H. E., and Goldberger, A. L. (1994). Mosaic organization of dna nucleotides. Phys Rev E Stat Phys Plasmas Fluids Relat Interdiscip Topics, 49:1685–1689.

Posner, M. I., Snyder, C. R., and Davidson, B. J. (1980). Attention and the detection of signals. Journal of Experimental Psychology: General, 109(2):160–174.

Reteig, L. C., Talsma, L. J., van Schouwenburg, M. R., and Slagter, H. A. (2017). Transcranial electrical stimulation as a tool to enhance attention. Journal of Cognitive Enhancement, 1:10–25.

Riddle, J., Hwang, K., Cellier, D., Dhanani, S., and D’Esposito, M. (2019). Causal Evidence for the Role of Neuronal Oscillations in Top–Down and Bottom–Up Attention. Journal of Cognitive Neuroscience, 31(5):768–779.

Romanella, S., Mencarelli, L., Burke, M., Rossi, S., Kaptchuk, T., and Santarnecchi, E. (2023). Targeting neural correlates of placebo effects. Cognitive, Affective, & Behavioral Neuroscience, 23:217–236.

Sadeghi, N. and Nazari, M. A. (2015). Effect of neurofeedback on visual-spatial attention in male children with reading disabilities: An event-related potential study. Neuroscience and Medicine, 6:71–79.

Saito, H., Kanayama, S., and Takahashi, T. (1992). Right angular lesion and selective impairment of motion vision in left visual field. The Tohoku Journal of Experimental Medicine, 166(2):229–238.

Schmidt, S. L., Iyengar, A. K., Foulser, A. A., Boyle, M. R., and Fröhlich, F. (2014). Endogenous cortical oscillations constrain neuromodulation by weak electric fields. Brain Stimulation, 7(6):878–889.

Seabold, S. and Perktold, J. (2010). statsmodels: Econometric and statistical modeling with python. In 9th Python in Science Conference.

Shapiro, S. S. and Wilk, M. B. (1965). An analysis of variance test for normality (complete samples). Biometrika, 52(3/4):591–611.

Slepian, D. (1978). Prolate spheroidal wave functions, fourier analysis, and uncertainty — v: the discrete case. The Bell System Technical Journal, 57(5):1371–1430.

Somers, D. C. and Sheremata, S. L. (2013). Attention maps in the brain. WIREs Cognitive Science, 4(4):327–340.

Sotero, R. C. (2016). Topology, cross-frequency, and same-frequency band interactions shape the generation of phase-amplitude coupling in a neural mass model of a cortical column. PLOS Computational Biology, 12(11):1–29.

Spooner, R. K., Wiesman, A. I., Proskovec, A. L., Heinrichs-Graham, E., and Wilson, T. W. (2020). Prefrontal theta modulates sensorimotor gamma networks during the reorienting of attention. Human Brain Mapping, 41(2):520–529.

Studer, B., Cen, D., and Walsh, V. (2014). The angular gyrus and visuospatial attention in decision-making under risk. NeuroImage, 103:75–80.

Sugimura, K., Iwasa, Y., Kobayashi, R., Honda, T., Hashimoto, J., Kashihara, S., Zhu, J., Yamamoto, K., Kawahara, T., Anno, M., Nakagawa, R., Hatano, K., and Nakao, T. (2021). Association between long-range temporal correlations in intrinsic eeg activity and subjective sense of identity. Scientific Reports, 11(1):422.

Sur, S. and Sinha, V. K. (2009). Event-related potential: An overview. Industrial Psychiatry Journal, 18(1).

Szewczyk, M., Augustynowicz, P., and Szubielska, M. (2022). Implicit spatial sequential learning facilitates attentional selection in covert visual search. an event-related potentials study. Frontiers in Human Neuroscience, 16.

ten Oever, S., de Graaf, T. A., Bonnemayer, C., Ronner, J., Sack, A. T., and Riecke, L. (2016). Stimulus presentation at specific neuronal oscillatory phases experimentally controlled with tacs: Implementation and applications. Frontiers in Cellular Neuroscience, 10.

Thut, G., Nietzel, A., Brandt, S. A., and Pascual-Leone, A. (2006). α-band electroencephalographic activity over occipital cortex indexes visuospatial attention bias and predicts visual target detection. Journal of Neuroscience, 26(37):9494–9502.

Veniero, D., Vossen, A., Gross, J., and Thut, G. (2015). Lasting eeg/meg aftereffects of rhythmic transcranial brain stimulation: Level of control over oscillatory network activity. Frontiers in Cellular Neuroscience, 9.

Vergara, V. M., Weiland, B. J., Hutchison, K. E., and Calhoun, V. D. (2018). The impact of combinations of alcohol, nicotine, and cannabis on dynamic brain connectivity. Neuropsychopharmacology, 43:877– 890.

Virtanen, P., Gommers, R., Oliphant, T. E., Haberland, M., Reddy, T., Cournapeau, D., Burovski, E., Peterson, P., Weckesser, W., Bright, J., van der Walt, S. J., Brett, M., Wilson, J., Millman, K. J., Mayorov, N., Nelson, A. R. J., Jones, E., Kern, R., Larson, E., Carey, C. J., Polat, ?I., Feng, Y., Moore, E. W., VanderPlas, J., Laxalde, D., Perktold, J., Cimrman, R., Henriksen, I., Quintero, E. A., Harris, C. R., Archibald, A. M., Ribeiro, A. H., Pedregosa, F., van Mulbregt, P., and SciPy 1.0 Contributors (2020). SciPy 1.0: Fundamental Algorithms for Scientific Computing in Python. Nature Methods, 17:261–272.

Vöröslakos, M., Takeuchi, Y., Brinyiczki, K., Zombori, T., Oliva, A., Fernández-Ruiz, A., Kozák, G., Kincses, Z. T., Iványi, B., Buzsáki, G., and Berényi, A. (2018). Direct effects of transcranial electric stimulation on brain circuits in rats and humans. Nature Communications, 9(1):483.

Vossen, A., Gross, J., and Thut, G. (2015). Alpha power increase after transcranial alternating current stimulation at alpha frequency (α-tacs) reflects plastic changes rather than entrainment. Brain Stimulation, 8(3):499–508.

Wischnewski, M. and Schutter, D. J. (2017). After-effects of transcranial alternating current stimulation on evoked delta and theta power. Clinical Neurophysiology, 128(11):2227–2232.

Zaehle, T., Rach, S., and Herrmann, C. S. (2010). Transcranial alternating current stimulation enhances individual alpha activity in human eeg. PLOS ONE, 5(11):1–7.

Zani, A., Tumminelli, C., and Proverbio, A. M. (2020). Electroencephalogram (eeg) alpha power as a marker of visuospatial attention orienting and suppression in normoxia and hypoxia. an exploratory study. Brain Sciences, 10(3).

Zhang, G., Garrett, D. R., and Luck, S. J. (2023). Optimal filters for erp research i: A general approach for selecting filter settings. bioRxiv.

